# Clonal seeds in hybrid rice using CRISPR/Cas9

**DOI:** 10.1101/496042

**Authors:** Chun Wang, Qing Liu, Yi Shen, Yufeng Hua, Junjie Wang, Jianrong Lin, Mingguo Wu, Tingting Sun, Zhukuan Cheng, Raphael Mercier, Kejian Wang

## Abstract

Heterosis, the observation that first generation hybrids outcompete the parental lines, is widely used in increasing the productivity and yield of agricultural crops^1,2^. However, heterosis is lost in the following generations because of genetic segregation. In addition, the high cost of hybrid seed production hinders the application of heterosis in many crops. Clonal reproduction through seeds could be revolutionary for agriculture by allowing self-propagation of F_1_ hybrids^3,4^. Here we show that heterozygosity of F_1_ hybrid rice can be fixed and thus propagated without additional crossing. First, we showed that multiplex editing of three key meiotic genes^5,6^ in hybrid rice leads to the production of clonal diploid gametes and tetraploid seeds. Next, editing of the *MATRILINEAL* (*MTL*) gene that involved in fertilization^7,8^ results in the induction of haploid seeds in hybrid rice. By simultaneous editing of these four endogenous genes in hybrid rice using the CRISPR/Cas9 system, we obtained in one generation plants able to propagate clonally through seeds. This opens the possibility to fix heterozygosity of hybrid varieties in food crops.

---

Heterosis (also known as hybrid vigor) is a phenomenon whereby hybrid offspring of genetically diverse individuals display increased vigor relative to their homozygous parents. Heterosis has been widely applied in agriculture to dramatically improve the production and to broaden adaptability of crops^1,2^. However, the essential process of hybrid seed production increases the seed cost and even prohibits its application in many crops. It has been proposed to fix the heterosis of hybrid crop by introduction of apomixis^3^. Apomixis is an asexual reproductive strategy where the offspring were generated through seeds, but without meiosis and fertilization. Although it has been described in many flowering plant taxa^9^, apomixis has not been reported in major crops. Previously, it was revealed that combined mutations of three genes that affect key meiotic processes created a genotype called *MiMe* (*Mitosis instead of Meiosis*) in which meiosis is totally replaced by mitotic-like division, leading to the production of male and female clonal diploid gametes in Arabidopsis and rice^5,6^. However, the self-fertilization of *MiMe* resulted in doubling of ploidy at each generation. By crossing Arabidopsis *MiMe* with CenH3-mediated chromosome elimination line, clonal diploid offspring were obtained^4^. However, the system still relies on the crossing between different plants and the CENH3-mediated chromosome elimination appeared to be difficult to transfer to other species^10^. Therefore, further work is required to achieve the aim of heterosis fixation in self-fertilized hybrids.

Firstly, to test the feasibility of *MiMe* technology in hybrid rice varieties, we performed experiments on Chunyou84 (CY84), an elite inter-subspecific hybrid rice from a cross between the maternal Chunjiang 16A (16A), a *japonica* male sterile line, and the paternal C84, an *indica-japonica* intermediate type line (Extended Data Fig.1). To ensure rapid generation of *MiMe* in the hybrid CY84 background, we simultaneously edited the *REC8*, *PAIR1* and *OSD1* genes using our previously developed multiplex CRISPR/Cas9 system^11^ (Fig. 1a). In the primary transformed plants, 7 of 32 plants were identified as frameshift triple mutants, and three of them were analyzed (Extended Data Fig.2). The triple mutant (*MiMe*) could not be distinguished from the wild-type CY84 based on its growth or morphology (Extended Data Fig.3). To test whether the meiosis was turned into a mitotic-like division, we investigated the male meiotic chromosome behavior in both wild type and *MiMe*. In the wild-type CY84 (Extended Data Fig.4a-f), 12 bivalents were scattered at diakinesis and aligned along the equatorial plate at metaphase I. The 12 pairs of homologous chromosomes separated at anaphase I and produced tetrad spores after the second meiotic division. In *MiMe* (Extended Data Fig.4g-i), 24 univalents were found in diakinesis and aligned at metaphase I. In anaphase I, 24 pairs of chromatids segregated into two groups and produced dyads of spores, suggesting that the meiosis has been turned into a mitotic-like division. We next examined the ploidy of spores of *MiMe* by performing fluorescent in situ hybridization (FISH) analyses using a 5S rDNA-specific probe, which identifies chromosome 11 of rice. Only one signal was observed in CY84 spores (n=30), while two signals were constantly observed in *MiMe* spores (n=40, Fig. 1b), showing that diploid gametes were generated in *MiMe*. We also investigated the fertility of *MiMe* mutant and found that the panicle seed setting rate in *MiMe* was ~81.2% (n=4043), which is comparable to that of wild type (~79.1%, n=3876), (Fig.1c, Table 1), suggesting that simultaneously editing of these three genes do not obviously affect fertility in this hybrid variety. The ploidy of the progeny of *MiMe* plant was investigated by flow cytometry and all (n=123) were found to be tetraploid plants (Fig.1d, Table 1). Further, we found that these progenies (n=123) retained completely the heterozygosity of their parent CY84 for 10 tested Insertion-deletion (Indel) makers (Fig.1e). And these progenies of *MiMe* displayed reduced fertility, increased grain size and elongated awn length compared to wild type, all of which being typical characteristics of tetraploid rice (Fig.1f). These results show that the *MiMe* phenotype can be rapidly introduced into hybrid rice varieties using CRISPR/Cas9 genome editing technique.

**Figure 1.**
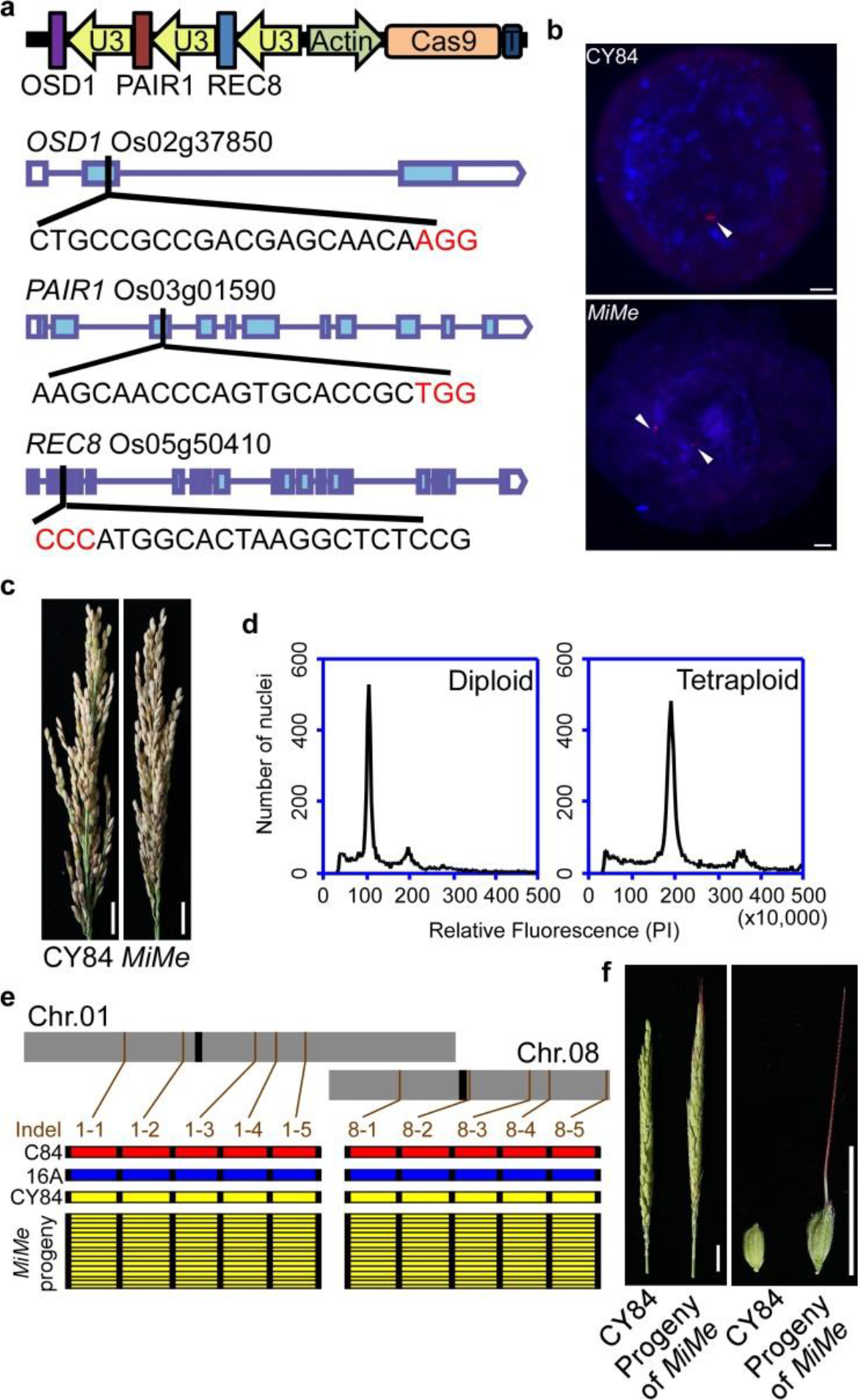
Turning meiosis into mitosis in hybrid rice variety Chunyou84 (CY84) **a**, Schematic diagram of the structure of CRISPR/Cas9 vector targeting *OSD1*, *PAIR1* and *REC8*. **b**, The chromosomes of CY84 and *MiMe* were probed by digoxige-nin-16-dUTP-labled 5S rDNA (red signal, indicated with white arrow) in spores, showing one signal in wild-type CY84 and two signals in *MiMe*. Scale bars, 5 μm. **c**, Panicles of wild-type CY84 and *MiMe*. The fertility of *MiMe* is as high as that of wild-type CY84. Scale bars, 2 cm. **d**, Ploidy analysis of CY84 (left) and the progeny of *MiMe* (right) by flow cytometry, which is found to be diploid and tetraploid, respectively (Table 1). **e**, Genotype analysis of the paternal C84, maternal Chunjiang 16A (16A), hybrid variety Chunyou84 (CY84) and the progeny siblings of *MiMe*. 10 Indel markers distributed on chromosomes 1 and 8 were used to identify the genotype of the offspring of *MiMe*. Positions of markers (brown) and centromeres (black) are indicated along the chromosomes. For each marker, plants carrying the C84 allele are in red, plants carrying the 16A allele are in blue, while plants with both C84 and 16A alleles appear in yellow. Each row represents one plant, and each column indicates a locus. **f**, Panicles and grain shape of CY84 and the progeny of *MiMe*. The progeny of *MiMe* displayed reduced fertility, increased glume size and elongated awn length. Scale bars, 2 cm.

**Table1.** Ploidy analysis of the progeny of CY84, *MiMe*, *mtl* and *Fix* lines.

|  | Line | Seed setting percentage | Progeny tested | Haploid+DH (%) | Diploid (%) | Tetraploid (%) |
| --- | --- | --- | --- | --- | --- | --- |
| CY84 | #1 | 77.2% (1151/1490) | 65 | 0 | 65 | 0 |
|  | #2 | 81.3% (951/1170) | 73 | 0 | 73 | 0 |
|  | #3 | 79.1% (962/1216) | 82 | 0 | 82 | 0 |
| *MiMe* | #1 | 81.9% (1178/1439) | 35 | 0 | 0 | 35 (100%) |
|  | #2 | 79.2% (877/1108) | 43 | 0 | 0 | 43 (100%) |
|  | #3 | 82.1% (1228/1496) | 45 | 0 | 0 | 45 (100%) |
| *mtl* | #1 | 9.1% (101/1103) | 77 | 6+0 (7.8%) | 71 | 0 |
|  | #2 | 13.6% (217/1601) | 90 | 2+1 (3.3%) | 87 | 0 |
|  | #3 | 11.3% (280/2476) | 81 | 1+1 (2.5%) | 79 | 0 |
| *Fix* | #1 | 3.7% (63/1725) | 39 | 0 | 2 (5.1%) | 37 |
|  | #2 | 5.2% (124/2373) | 64 | 0 | 3 (4.7%) | 61 |
|  | #3 | 4.3% (76/1752) | 42 | 0 | 4 (9.5%) | 38 |

*MiMe* clonal gametes participate in normal self-fertilization, giving rise to progeny with doubled ploidy. This ploidy doubling must be prevented to achieve apoximis. Recently, it was reported that the *MATRILINEAL* (*MTL*) gene, a sperm-specific phospholipase, triggers haploid induction in maize^7,8^. To test whether the homologous gene could be manipulated to induce haploid in self-fertilized hybrid rice, we edited the *MTL* gene in CY84 (Fig. 2a). 11 of 32 transformed plants were identified as frameshift mutants, and three of them were analyzed (Extended Data Fig.5). The *mtl* mutants showed normal vegetative growth (Extended Data Fig.3), but the seed-setting rates significantly reduced to ~11.5% (n=5180, Fig. 2b, Table 1). 12 Indel markers (1 per chromosome) that were polymorphic between the two parents were used to determine the genotype of the progenies of *mtl* plants (Extended Data Table1). In the wild-type CY84 progeny, no plants homozygous at all markers were found (n=220, Table 1). In contrast, 11 plants among 248 *mtl* progenies appeared to be homozygous for all markers (Fig. 2c, Table 1). Flow cytometry results showed that 9 of these plants were indeed haploid, while 2 were diploid, presumably resulting from spontaneous doubling of haploid embryos (Fig. 2d, Table 1). To further classify the genotype of those identified plants, the whole genomes of 2 haploids, 2 doubled haploids of *mtl* progenies, and 2 offspring plants of wild-type CY84 were resequenced with a depth of 30-fold. A total of 78,909 single nucleotide polymorphisms (SNPs) that differed between two parents were screened out for detailed genotype analysis. Whole genome sequencing revealed that the haploids and doubled haploids were homozygous at all loci (Fig. 2e), and recombinant compared to the parental genome, suggesting that they are respectively derived from a single gamete. The haploid plants showed reduced plant height, decreased glume size and loss of fertility, while the doubled haploid plant displayed normal vegetative and reproductive growth (Fig. 2f). The results demonstrated that haploid plants can be generated by self-fertilization of hybrid varieties.

**Figure 2.**
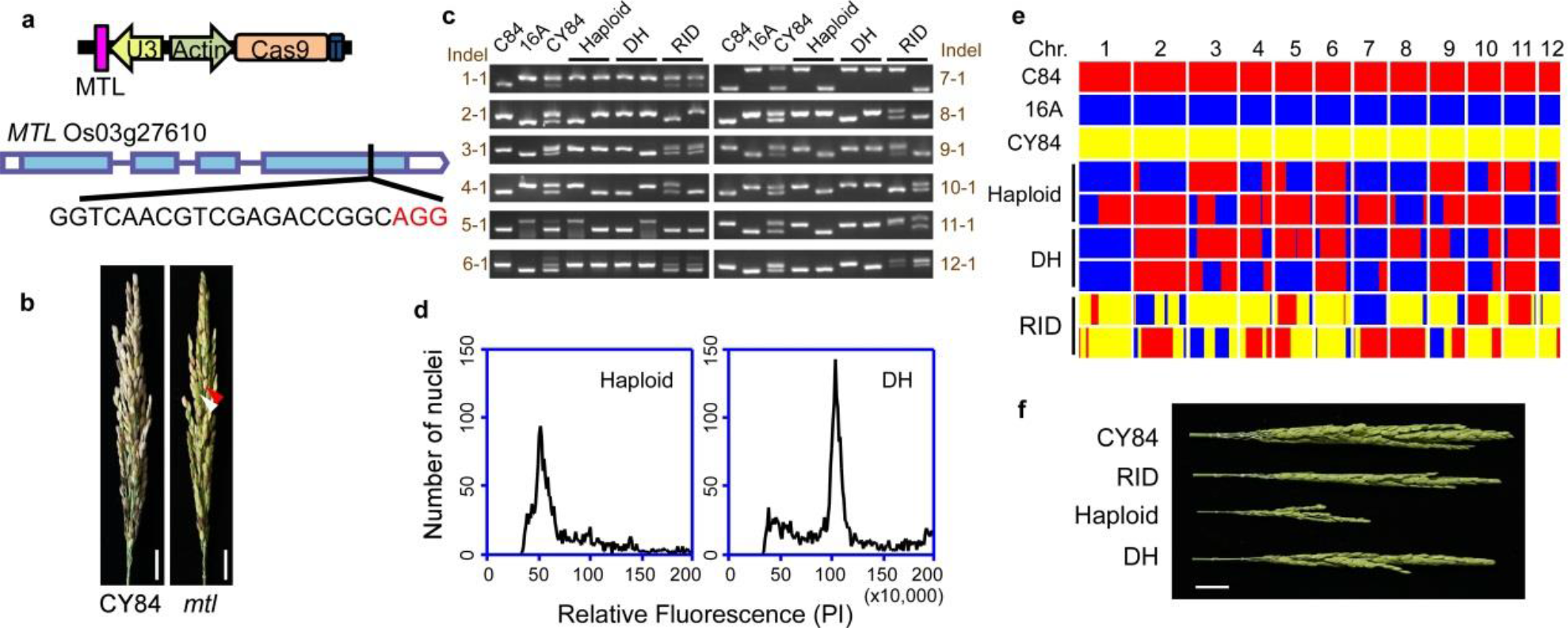
Generation of haploid inducer line by editing the *MTL* gene in hybrid rice variety CY84. **a**, Schematic diagram of the structure of CRISPR/Cas9 vector targeting *MTL*. **b**, Panicles of the WT and *mtl* in CY84 background. The fertility was decreased in *mtl*, white arrow indicates aborted seed, and red arrow shows fertile seed. Scale bars, 2 cm. **c**, Determination of the genotype of haploids, doubled haploids (DH) and recombinant inbred diploids (RID) using 12 Indel markers (1 per chromosome). Plants homozygous at all markers in the progeny siblings of *mtl* were identified as haploid or DH. **d**, Ploidy analysis of the haploid and DH by flow cytometry (Table 1). **e**, Whole genome sequencing of the haploid, DH and RID plants. 12 blocks represent 12 chromosomes. The SNPs of C84 allele are in red, the SNPs of 16A allele are in blue, and co-existence of both alleles are in yellow. **f**, Panicles of wild-type CY84 and *mtl* progeny, including RID, haploid and DH plants. Scale bars, 2 cm.

Since turning meiosis into mitosis and paternal genome elimination is possible in self-fertilized hybrid rice, we next test the possibility of inducing heterozygosity fixation without additional crossing in hybrid rice by simultaneously editing four genes, namely *OSD1*, *PAIR1*, *REC8* and *MTL* in CY84 (Fig. 3a-b). Among 22 transgenic plants, three were identified by DNA sequencing as *osd1/pair1/rec8/mtl* quadruple mutants (namely *Fix*, *<u>Fix</u>ation of hybrids*) and used for further analysis (Extended Data Fig.6). The *Fix* mutants grew normally during the vegetative stage (Fig. 3c). During reproductive stage, the male meiotic chromosome behavior was investigated and found to be indistinguishable from that of *MiMe* (Extended Data Fig.4j-l). The panicle seed setting percentage was found to be ~4.5% (n=5850) (Table 1, Fig. 3c), which is slightly lower than that of the *mtl* mutant. In the progeny seedlings, the ploidy was investigated using flow cytometry. Among 145 progeny of *Fix* mutants, 136 were identified as tetraploid and 9 as diploid (Fig. 3d, Table 1). To investigate whether the heterozygosity was fixed in these diploid offspring, the genomes of 2 diploid and 2 tetraploid offspring plants of *Fix* were resequenced with an average of 30× coverage. Bioinformatic analysis revealed that all the 78,909 SNPs were heterozygous in both these diploid and tetraploid progeny plants, and were thus genetically identical to the hybrid rice CY84 (Fig. 3e). Finally, we investigated the phenotype of the potential clonal plants of *Fix*. All these 9 diploid plants displayed normal glume size and awn length, and showed a reduced seed setting (~10%, n=2726), which were similar to their parent *Fix* plants (Fig. 3f). Taken together, the diploid progeny of *Fix* plant displayed the same ploidy, the same heterozygous genotype, and the similar phenotype with the parent *Fix* plants, implying that *Fix* is able to produce clonal seeds and fix the heterozygosity of hybrid rice.

**Figure 3.**
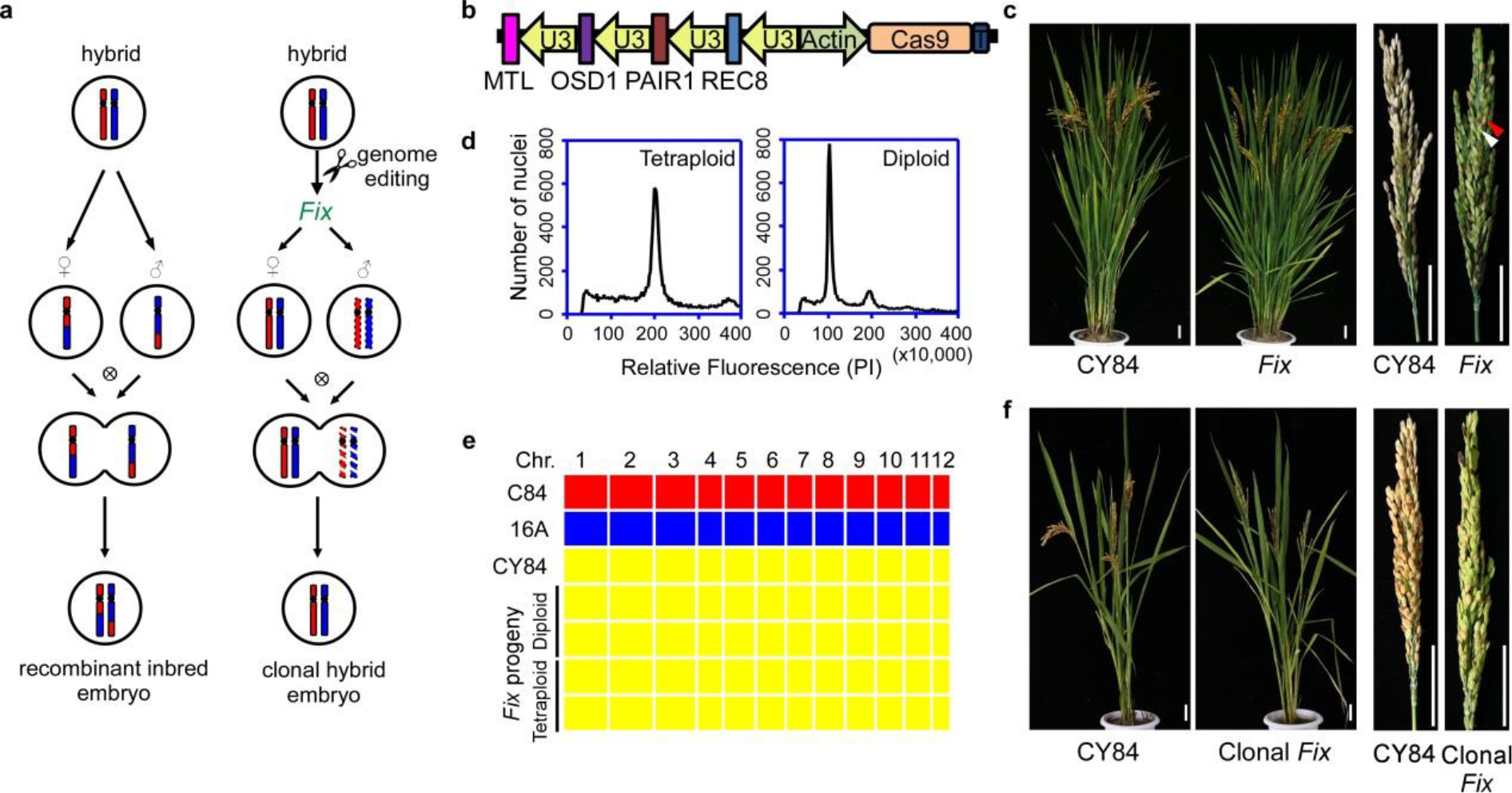
Fixation of rice heterozygosity by multiplex gene editing in hybrid rice variety CY84. **a**, The model of fixation of heterozygosity of hybrid. In normal sexual reproduction (left), recombinant inbred embryos are generated by fusion of recombined haploid gametes. The clonal reproduction strategy (right) is based on two components: meiosis is turned into mitosis to produce clonal diploid gametes (*MiMe*), and the genome of male gamete is eliminated by knocking out the *MTL* gene. The progeny of self-fertilized *Fix* is genetically identical to the hybrid parent. **b**, Schematic diagram of the structure of CRISPR/Cas9 vector simultaneously targeting *OSD1*, *PAIR1*, *REC8* and *MTL*. **c**, Comparison of the morphology and panicles of CY84 and *Fix* (*osd1 pair1 rec8 mtl*). The fertility was decreased in *Fix*. An aborted seed is indicated with white arrow, and a normally developed seed is indicated with red arrow. Scale bars, 5 cm. **d**, Ploidy analysis of the progeny of *Fix* by flow cytometry, including tetraploid (left) and diploid (right), respectively. **e**, Whole genome sequencing of the diploid and tetraploid progenies of *Fix*. The SNPs of C84 allele are in red, the SNPs of 16A allele are in blue, and co-existence of both alleles are in yellow. 12 blocks represent 12 chromosomes. The diploid and tetraploid progenies of *Fix* are heterozygous, identical to CY84. **f**, Comparison the morphology and panicles of wild-type CY84 and the diploid progeny of *Fix*. Both plants were grown in the glasshouse. The clonal *Fix* displayed normal growth except the reduced fertility, which is similar to that of parent *Fix* plant. Scale bars, 5 cm.

Our findings revealed that hybrids can be self-propagated through seeds by targeted editing of four endogenous genes in rice hybrid varieties. Simultaneous editing of *REC8*, *PAIR1* and *OSD1* genes does not have obvious adverse effects on the growth and reproduction of the hybrid. On contrast, the *MTL* gene used to induce paternal genome elimination has impacts on rice fertility and is not fully penetrant; further work is thus required to allow this technology to reach the rice fields. However, the findings in this study revealed a strategy to fix heterozygosity in rice. Considering the establishment of multiplex genome editing technology in many other crops along with the conservation of these four genes, the strategy might extend heterosis application in agriculture.

## Methods

### Plasmid construction

The plasmids expressing the CRISPR/Cas9 system were constructed *via* the isocaudamer ligation method, as previously described^11^. The modified single guide RNAs (sgRNAs) scaffold and *ACTIN1* promoter-driven Cas9 were used to increase the mutation rate in this study^12^. Briefly, the double-stranded overhangs of target oligoes (listed in Extended Data Table1) were ligated into the SK-sgRNA vectors digested with *Aar*I. Then the sgRNAs of *OSD1* (digested with *Kpn*I and *Sal*I), *PAIR1* (digested with *Xho*I and *Bgl*II) and *REC8* (digested with *Bam*HI and *Nhe*I) were assembled into one pC1300-ACT:Cas9 binary vector (digested with *Kpn*I and *Xba*I) using T4 ligase to obtain the vector pC1300-ACT:Cas9-sgRNA^OSD1^-sgRNA^PAIR1^ -sgRNA^REC8^ for generation of *MiMe*. The sgRNA of *MTL* (digested with *Kpn*I and *Nhe*I) was assembled into pC1300-ACT:Cas9 binary vector (digested with *Kpn*I and *Xba*I) to obtain the vector pC1300-ACT:Cas9-sgRNA^MTL^ for generation of *mtl*. The sgRNA of *MTL* (digested with *Kpn*I and *Nhe*I) was assembled into pC1300-ACT:Cas9-sgRNA^OSD1^-sgRNA^PAIR1^-sgRNA^REC8^ vector (digested with *Kpn*I and *Xba*I) to obtain the vector pC1300-ACT:Cas9-sgRNA^OSD1^-sgRNA^PAIR1^-sgRNA^REC8^-sgRNA^MTL^ for generation of *Fix*.

### Rice transformation and growth conditions

The hybrid rice Chunyou 84 (CY84) was used as the host variety in this study. The generation of transgenic rice, by *Agrobacterium*-mediated transformation with the strain EHA105, was performed by the Biogle company (Hangzhou, China).

The T_0_ generation of transgenic plants were grown in the transgenic paddy fields of the China National Rice Research Institute in Hangzhou, China (at N 30.32°, E 120.12°) in the summer of 2017. The T_1_ plants were grown in greenhouse in the winter of 2017.

### Detection of genome modifications

Genomic DNA was extracted from approximately 100 mg of rice leaf tissue *via* the CTAB method. PCR was conducted with KOD FX DNA Polymerase (Toyobo, Osaka, Japan) to amplify the genomic regions surrounding the target sites. The primers are listed in Extended Data Table1. The fragments were sequenced by the Sanger method and decoded by the degenerate sequence decoding method^13^.

### Cytological analyses

Young panicles of meiosis stage were harvested and fixed in Carnoy’s solution (ethanol:glacial acetic, 3:1). Microsporocytes undergoing meiosis were squashed in an acetocarmine solution. Slides were frozen in liquid nitrogen and the coverslips were removed with a blade quickly. Chromosomes were counterstained with 4’,6-diamidinophenylindole (DAPI) in an antifade solution (Vector Laboratories, Burlingame, CA). Microscopy was conducted using an Olympus BX61 fluorescence microscope with a microCCD camera.

Fluorescence *in situ* hybridizaiton (FISH) analysis was conducted as described previously^14^. The plasmid pTa794 was used as FISH probe to quantify the 5S rDNA.

### Genotyping with Indel Markers

Insertion-deletion (Indel) markers to distinguish genotypes of heterozygote and homozygote were designed based on the whole-genome sequences of C84 and 16A. The primers are listed in Extended Data

Table1. The genotyping was performed by normal PCR program using 2× Taq Master Mix (Novoprotein Scientific, China), and the PCR products were detected using 5% agarose gels.

### Flow cytometry determination of DNA content in leaf cell nuclei

The ploidy of leaf cell was determined by estimating nuclear DNA content using flow cytometry. All procedures were done at 4 °C or on ice. Approximately ~ 2 cm^2^ of leaf tissue was chopped using a new razor blade for 2 to 3 minutes in 1 ml LB01 Buffer (15 mM Tris, 2 mM Na_2_EDTA, 0.5 mM spermine tetrahydrochloride, 80 mM KCl, 20 mM NaCl, 0.1% Triton X-100, 15 mM β-mercaptoethanol, pH 7.5, filter through a 0.22 μm filter). The homogenate was filtered through the 40-μm nylon filter followed by centrifugation (1200× rpm, 5 min) to collect the nuclei. The supernatant was discarded and the pellet was resuspended with 450 μL of fresh LB01 Buffer, then 25 μl of 1 mg/ml propidium iodide (PI, Sigma P4170) and 25 μl of 1 mg/ml DNase-free RNase A (Sigma V900498) were added to stain the DNA. The stained samples were incubated on ice in darkness for 10 minutes prior to analysis. The samples were analyzed using BD Accuri C6 flow cytometer, with the laser illumination at 552 nm and 610/20 nm filter. The gating strategy was provided in Supplementary Information. Samples with the same result of CY84 were deemed as diploids, which the first peak of relative fluorescence at ~100 (x10,000). And the samples with the first peak of relative fluorescence at ~50 (x10,000) were deemed as haploids, while samples with the first peak of relative fluorescence at ~200 (x10,000) were deemed as tetraploids.

### Whole genome re-sequencing and genotype calling

The 150-bp paired-end reads were generated by Illumina Hiseq2500, covering approximately an average depth of 30× for each sample. The short-read sequence data have been deposited in the NCBI Sequence Read Archive (SRP149641, SRP149677). The raw paired-end reads were first filtered into clean data using NGSQCtookit v2.3.3^15^. The cutoff value for PHRED quality score was set to 30. Clean reads of each accession were aligned against the rice reference genome (IRGSP 1.0) using the software SOAPaligner (soap version 2.21) ^16^ with the parameters of ‘-m 200, -x 1000, -l 35, -s 42, -v 5’ and ‘−p 8’. To get high-quality SNPs, reads that could be mapped to different genomic positions were excluded by SOAPsnp^17^. Uniquely mapped single-end and paired-end results were used in the SNP calling. Genotype calling was carried out in the whole genome region using these SNPs which are heterozygous in the parent. The window size (the number of n consecutive SNPs in a window) was 0.1 K. And the recombination map was constructed for each chromosome.

## Data availability

Whole genome sequencing data are deposited in the NCBI Sequence Read Archive (SRP149641, SRP149677). Patent applications have been filed relating to work in this manuscript.

## Supporting information

## Acknowledgements

This research was supported by the Agricultural Science and Technology Innovation Program of Chinese Academy of Agricultural Sciences, and the National Natural Science Foundation of China (No. 31401363).

## Author Contributions

C.W., and K.W. conceived and designed the study. C.W., Y.S., and Z.C. performed the lab experiments. Q.L., and T.S. conducted the computational analyses. Y.H., and J.W. carried out the field experiments. J.L., and M.W. provided the rice varieties and helped with the field management. C.W., R.M., and K.W. wrote the manuscript.

