## Supplementary material for "Clonal seeds in hybrid rice using CRISPR/Cas9"

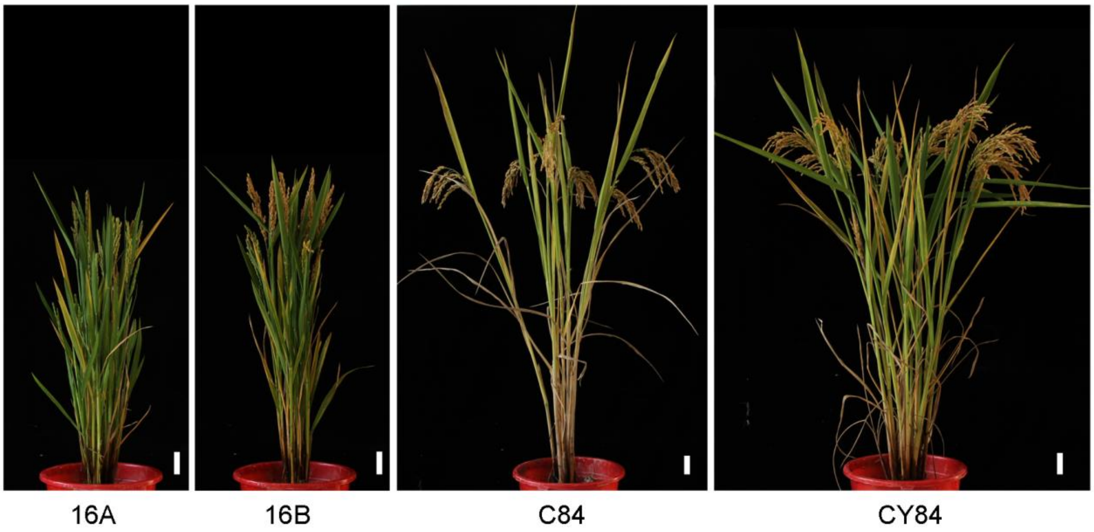


**Extended Data Figure 1 | The morphology of inter-subspecific hybrid rice Chunyou84 (CY84) and its parents.** Chunjiang 16A (16A), a late *japonica* male sterile line. C84, an *indica-japonica* intermediate type restore line with wide compatibility. The maintainer line Chunjiang 16B (16B) is used to maintain the sterility of 16A. Scale bars, 5 cm.

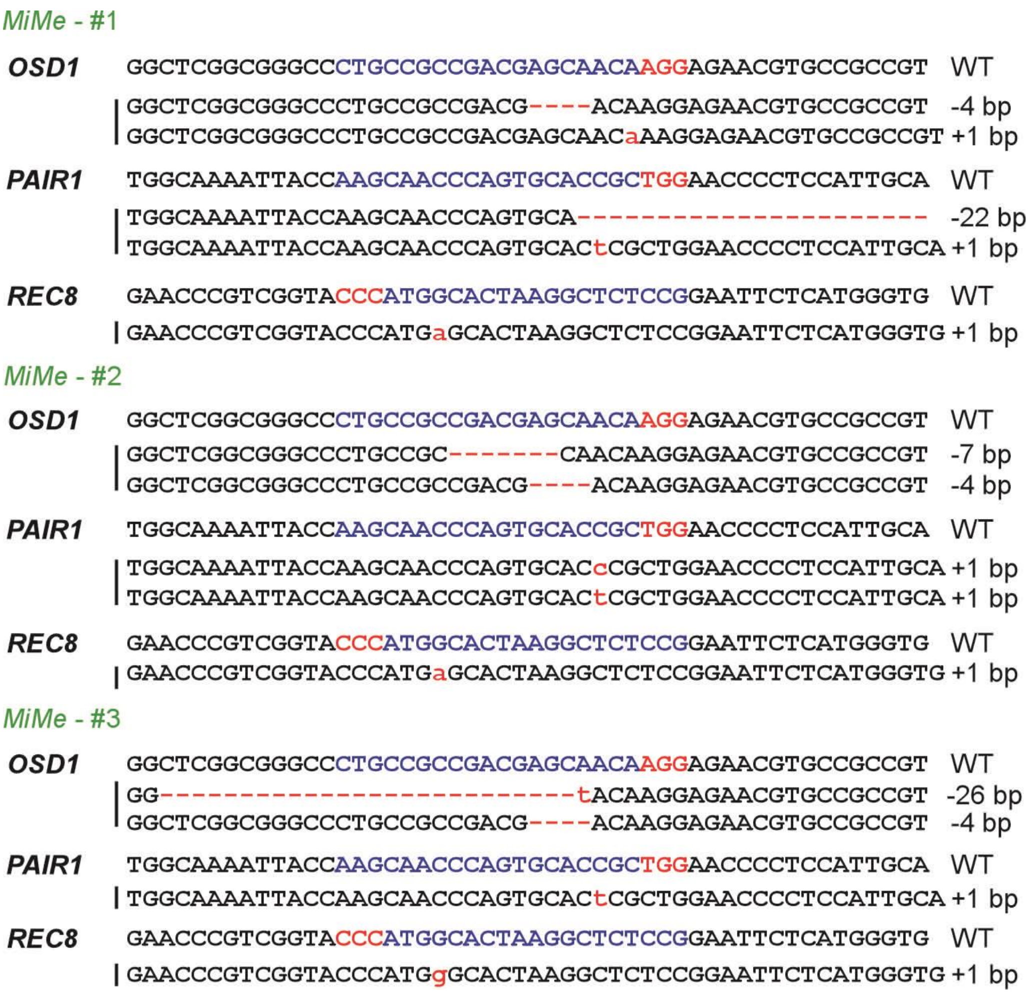


**Extended Data Figure 2| Sequencing assay of mutations around the targets of *OSD1*, *PAIR1* and *REC8* in *MiMe*.** The PAM and target sequence of wild type (WT) are denoted in red and blue, respectively. - indicates deleted nucleotides.

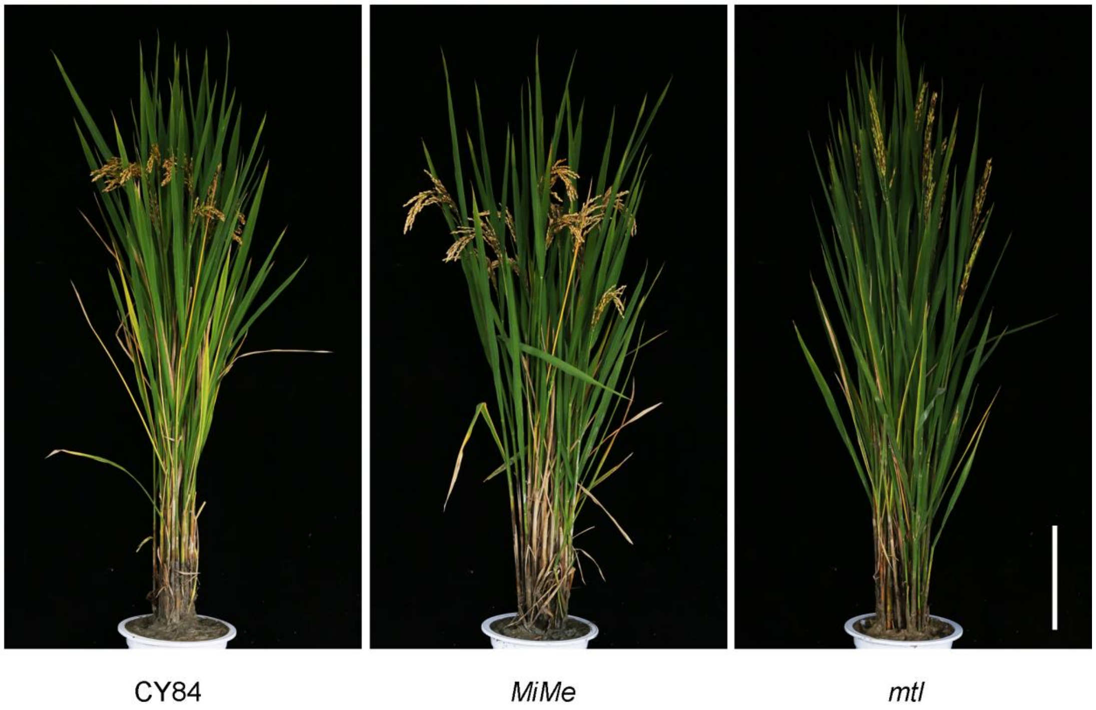


**Extended Data Figure 3| The morphology of wild-type CY84, *MiMe* and *mtl*.** The *MiMe* and *mtl* mutants showed normal vegetative growth. Scale bars, 20 cm.

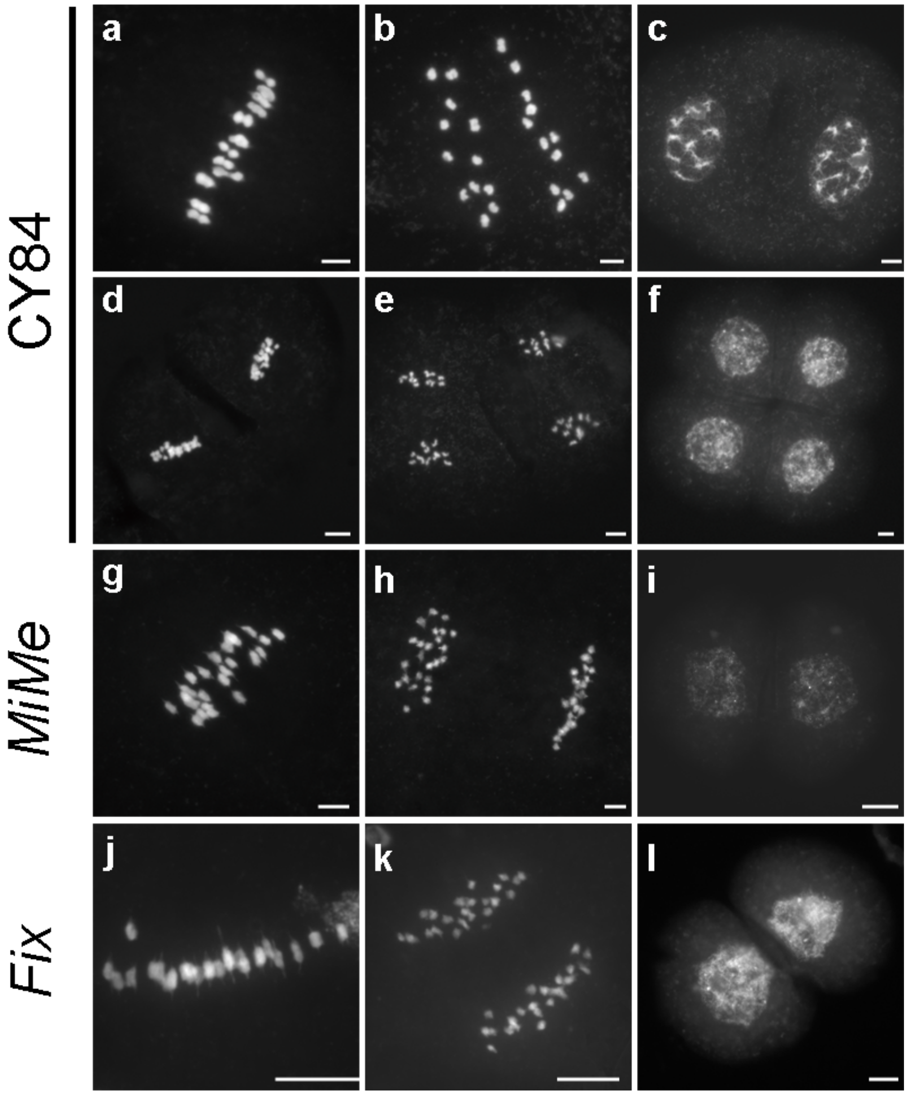


**Extended Data Figure 4 | Chromosome spreads of male meiosis in wild-type CY84, *MiMe* and *Fix*. a-f**, CY84 (n=32). **a**, Metaphase I with 12 aligned bivalents. **b**, Anaphase I. **c**, Telophase I. **d**, Metaphase II. **e**, Anaphase II. **f**, Telophase II. **g-i**, *MiMe* (n=45). **g**, Metaphase I with 24 aligned univalents. **h**, Anaphase I with segregation of 24 pairs of chromatids. **i**, Telophase I. **j-l**, *Fix* (n=52). **j**, Metaphase I with 24 aligned univalents. **k**, Anaphase I with segregation of 24 pairs of chromatids. **i**, Telophase I. Scale bars, 5 μm.

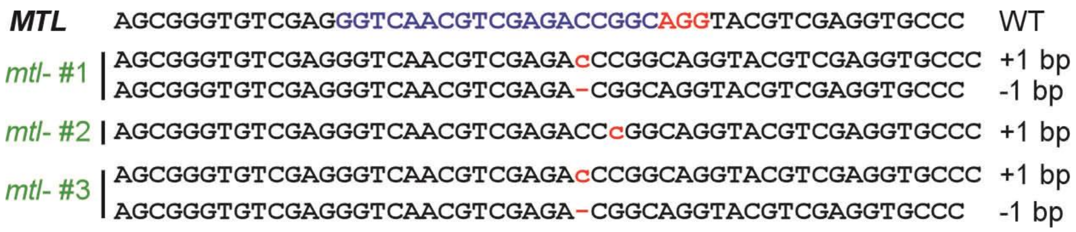


**Extended Data Figure 5 |Sequencing assay of mutations around the target of *MTL*.** The PAM and target sequence of wild type (WT) are denoted in red and blue, respectively. - indicates deleted nucleotides.

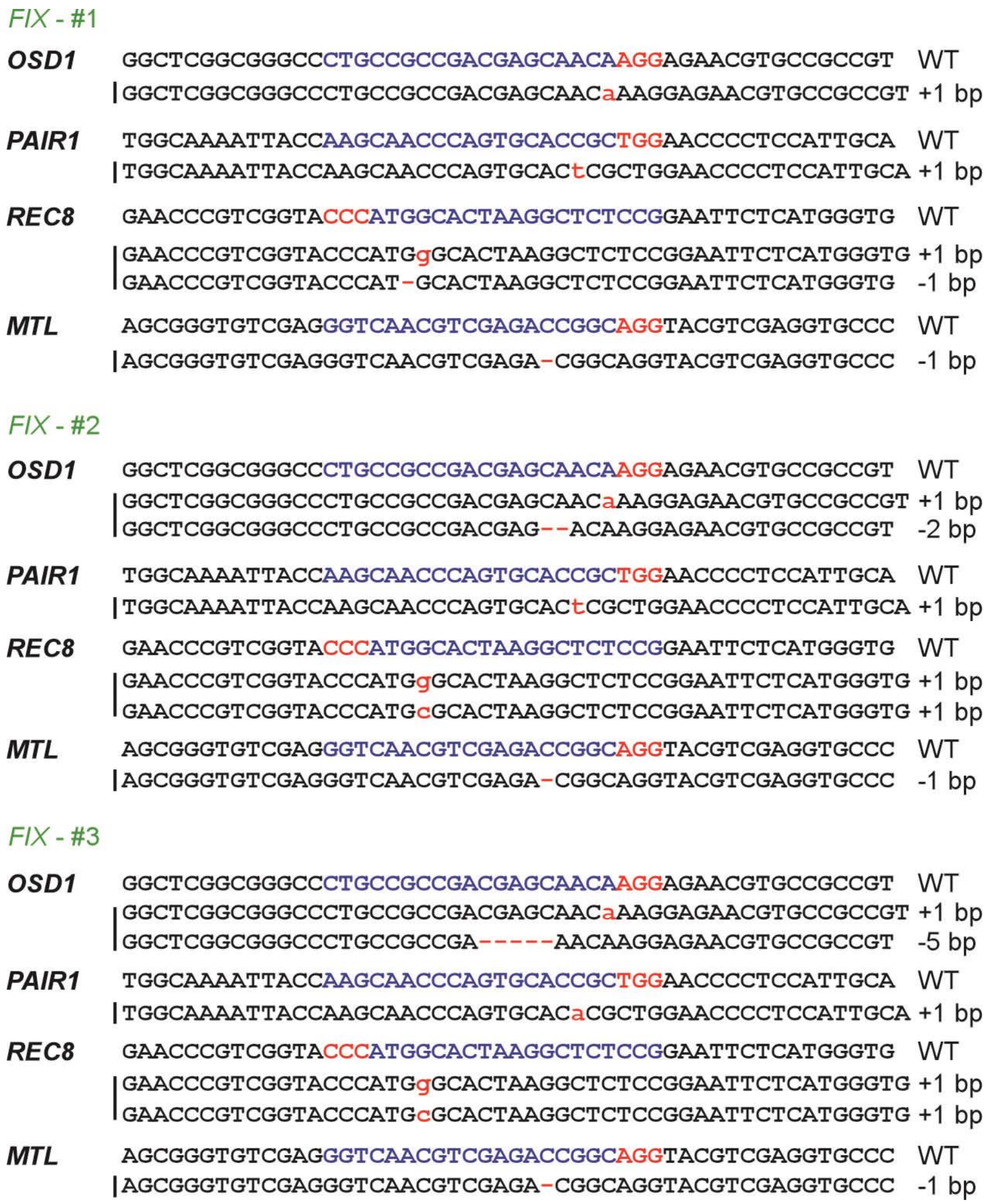


**Extended Data Figure 6 |Sequencing assay of mutations around the targets of *OSD1*, *PAIR1*, *REC8* and *MTL* in *Fix*.** The PAM and target sequence of wild type (WT) are denoted in red and blue, respectively. - indicates deleted nucleotides.

### Extended DataTable1|Primers used in this study.

| Primer | Sequence(5'-3') | Remarks |
|---|---|---|
| OSD1-g++ | ggcaCTGCCGCCGACGAGCAACA | OSD1 target oligo of sgRNA |
| OSD1-g-- | aaacTGTTGCTCGTCGGCGGCAG | OSD1 target oligo of sgRNA |
| PAIR1-g++ | ggcaAAGCAACCCAGTGCACCGC | PAIR1 target oligo of sgRNA |
| PAIR1-g-- | aaacGCGGTGCACTGGGTTGCTT | PAIR1 target oligo of sgRNA |
| REC8-g++ | ggcaCGGAGAGCCTTAGTGCCAT | REC8 target oligo of sgRNA |
| REC8-g-- | aaacATGGCACTAAGGCTCTCCG | REC8 target oligo of sgRNA |
| MTL-g++ | ggcaGGTCAACGTCGAGACCGGC | MTL target oligo of sgRNA |
| MTL-g-- | aaacGCCGGTCTCGACGTTGACC | MTL target oligo of sgRNA |
| OSD1-F | ATCTCCAGGATGCCTGAAGTGAG | primer for detection of genome modification |
| OSD1-R | CCTAGACTGCTACTCTTGCTAGTGAT | primer for detection of genome modification |
| PAIR1-F | CTGTACCTGTGCATCTAATTACAG | primer for detection of genome modification |
| PAIR1-R | CCCCATCTTATGTACTGAGCTTGCCAG | primer for detection of genome modification |
| REC8-F | GCGACGCTTCACTCGAAGATCA | primer for detection of genome modification |
| REC8-R | CGCCATGCCTCGTTGATCTCAA | primer for detection of genome modification |
| MTL-F | ACAGTGACTAGTGACAAACGATCG | primer for detection of genome modification |
| MTL-R | GATCGCGTCAGCATGATGCGTGTAC | primer for detection of genome modification |
| Indel 1-1-F | GTTATTTCACCCGGGTCT | Indel marker primer, on chromosomes 1 |
| Indel 1-1-R | GCTCTAACCGTGTCATTCT | Indel marker primer, on chromosomes 1 |
| Indel 1-2-F | TGCTCTAGGCTGGGTTTAGT | Indel marker primer, on chromosomes 1 |
| Indel 1-2-R | CTAACACACCGAAGCATCAC | Indel marker primer, on chromosomes 1 |
| Indel 1-3-F | ATTACAGGGATGCACTGCTGAC | Indel marker primer, on chromosomes 1 |
| Indel 1-3-R | GAAGCCACTCTGAAATCGGCA | Indel marker primer, on chromosomes 1 |
| Indel 1-4-F | GTATCTCCCATCTAGATCTGC | Indel marker primer, on chromosomes 1 |
| Indel 1-4-R | ATTCCACATTGAGCGCTTGTC | Indel marker primer, on chromosomes 1 |
| Indel 1-5-F | AGCTCGCAATGGAGTCATCATG | Indel marker primer, on chromosomes 1 |
| Indel 1-5-R | TCTAATCCGTGTTTCCACTGC | Indel marker primer, on chromosomes 1 |
| Indel 2-1-F | CGTGGCCATCTTGTAGTG | Indel marker primer, on chromosomes 2 |
| Indel 2-1-R | GCTGCAGTAGACAGAGAT | Indel marker primer, on chromosomes 2 |
| Indel 3-1-F | CTTGCCCATCAGCCTATCAC | Indel marker primer, on chromosomes 3 |
| Indel 3-1-R | ACACGTACACAGCCATGAGA | Indel marker primer, on chromosomes 3 |
| Indel 4-1-F | TGCCTCTTTCGAACGTATCC | Indel marker primer, on chromosomes 4 |
| Indel 4-1-R | TAAGCTACGAGCAGTGGACA | Indel marker primer, on chromosomes 4 |
| Indel 5-1-F | GCGCACTACTTTTGAACCGT | Indel marker primer, on chromosomes 5 |
| Indel 5-1-R | GAGCGGTGAAAGGTAAAATT | Indel marker primer, on chromosomes 5 |
| Indel 6-1-F | ACGCCGCTAGGATATTGGAAGAC | Indel marker primer, on chromosomes 6 |
| Indel 6-1-R | TCCGACGCGGCACGAACCAACG | Indel marker primer, on chromosomes 6 |
| Indel 7-1-F | ATATGCACAAAGGTAGCGTG | Indel marker primer, on chromosomes 7 |
| Indel 7-1-R | TGCTATTATCGACAAGAAGG | Indel marker primer, on chromosomes 7 |
| Indel 8-1-F | AGCTTAATACGCAGAGCC | Indel marker primer, on chromosomes 8 |
| Indel 8-1-R | CATGGTCCTTCACGTGTA | Indel marker primer, on chromosomes 8 |

| | | |
|---|---|---|
| Indel 8-2-F | CAGGGAGTTGGAAACCTTTC | Indel marker primer, on chromosomes 8 |
| Indel 8-2-R | CCAAAATGCTAAACGGTGTG | Indel marker primer, on chromosomes 8 |
| Indel 8-3-F | AGGTCTTCTGTCCAAGTTCA | Indel marker primer, on chromosomes 8 |
| Indel 8-3-R | AACCATATAAACTCATCTGC | Indel marker primer, on chromosomes 8 |
| Indel 8-4-F | CATGCAGATAGCTCGCTTGT | Indel marker primer, on chromosomes 8 |
| Indel 8-4-R | CACCTCTCAGGACAACTGTA | Indel marker primer, on chromosomes 8 |
| Indel 8-5-F | TTGCTTGGAGTTTGACGACG | Indel marker primer, on chromosomes 8 |
| Indel 8-5-R | TGGTGTCGGCCTATGCTAAT | Indel marker primer, on chromosomes 8 |
| Indel 9-1-F | ATTCTTGTGAGGACGGGAGG | Indel marker primer, on chromosomes 9 |
| Indel 9-1-R | GAGAGGCGGTTACCATCTGC | Indel marker primer, on chromosomes 9 |
| Indel 10-1-F | GCGCATCCATGCATATCCAA | Indel marker primer, on chromosomes 10 |
| Indel 10-1-R | GACAAGGTGTTGCCCAAGAA | Indel marker primer, on chromosomes 10 |
| Indel 11-1-F | GGCATCATTAAGGCTTGT | Indel marker primer, on chromosomes 11 |
| Indel 11-1-R | CTGGCGATCTCTGTGAGG | Indel marker primer, on chromosomes 11 |
| Indel 12-1-F | TTGACCTTACTAATCCGAGC | Indel marker primer, on chromosomes 12 |
| Indel 12-1-R | AGGTCGCGAGGTTTCTTGAT | Indel marker primer, on chromosomes 12 |
